# Attentional set to safety recruits the medial prefrontal cortex

**DOI:** 10.1101/249326

**Authors:** Shuxia Yao, Song Qi, Keith M. Kendrick, Dean Mobbs

**Affiliations:** The Clinical Hospital of Chengdu Brain Science Institute, MOE Key Laboratory for NeuroInformation, University of Electronic Science and Technology of China, Chengdu, Sichuan 611731, China; California Institute of Technology, Pasadena, California 91125; Columbia University in the City of New York, New York, NY 10027

**Author notes:** Corresponding authors: Dean Mobbs, Humanities and Social Sciences, 319 Baxter Hall, California Institute of Technology (Caltech), Pasadena, California, 91125. Keith M. Kendrick, No. 2006, Xiyuan Ave., West Hi-Tech Zone, Chengdu, Sichuan 611731, China.

## Abstract

During threat assessment, the early detection of danger is highly adaptive, yet the fast orientation towards safety is also key to survival. The present study aimed to explore how the human brain searches for safety by manipulating subjects’ attentional set to cues associated with shock probability. Subjects were asked to judge random dots motion (RDM) direction and could be shocked for incorrect responses (RDM task) while keeping alert in detecting the shock probability cues (cue detection task). In contrast to the safe condition, where subjects searched for cues associated with no shock probability, incorrect responses to ‘dangerous+’ (D+) cues would increase the shock probability and correct responses to ‘dangerous-’ (D-) cues would decrease shock probability. In the RDM task, results showed that relative to the D+, the safe attentional set resulted in stronger activation in the ventral medial prefrontal cortex (vmPFC), a core region involved in flexible threat assessment and safety signalling. The vmPFC was also recruited by the D-compared to the D + attentional set. In the cue detection task, shorter response times and greater accuracy were observed for D+ compared to D‐ and safe cues. Correspondingly, at the neural level D+ cues induced increased activity in the frontoparietal attention network including the inferior parietal lobule and intraparietal sulcus. Overall, our findings demonstrate that attentional set for searching safety recruits the vmPFC, while detection of threat elicits activity in the frontoparietal attention network, suggesting a new role for these regions in human defensive survival circuitry.

**Significance Statement:** While early detection of threat is highly adaptive, the fast orientation towards safety is also key to survival. However, little is known about neural mechanisms underlying attentional set to safety. Using a novel dots motion paradigm combined with fMRI, we explored how human brain prepares for safety searching by manipulating subjects’ attentional set to cues associated with shock probability. Relative to the dangerous attentional set associated with increasing shock probability, the safe attentional set resulted in stronger activity in the ventral medial prefrontal cortex, a core region involved in flexible threat assessment and safety signalling, suggesting a new role for this region in human defensive survival system in encoding stimuli with survival significance.

## Introduction

Our attention systems have evolved to detect stimuli that are of survival value, with early detection of potential ecological dangers being of crucial importance. During threat assessment, the detection of threat per se is only one among several other parallel strategies including threat monitoring, prediction and safety seeking (Mobbs et al., 2015). In the case of searching for safety, the organism will search the environment for a safe refuge and in turn, this will influence its decision to either freeze or flee. For example, it has been theorized that when escape is viable, flight will occur but when it is not then freezing will be the choice of defense (Blanchard and Blanchard, 1990). Consequently, decreased fear induced by the knowledge of being safe facilitates exploitation of the environment by organisms and thus increases their foraging and copulation opportunities (Cooper and Blumstein, 2015; Rogan et al., 2005). How the human brain implements this safety search strategy is unknown.

Safety searching is not only critical for increasing the probability of escape from predators but also affects fear perception. Animal studies have shown that in the presence of danger, safety cues can abolish innate defensive analgesia (Wiertelak et al., 1992). Fear conditioning studies have also provided evidence that learned safety (in the context of the unpaired neutral vs. the aversive conditioned stimulus) is associated mainly with the medial prefrontal cortex (mPFC) and basolateral amygdala (Likhtik, et al., 2014; Stujenske et al., 2014). Access to safety can decrease fear responses either in healthy populations (Grillon et al., 1994; Hood et al., 2010) or in patients with affective disorders such as panic disorder and claustrophobia (Carter et al., 1995; Telch et al., 1994). Human fMRI studies have further highlighted the role of the ventral mPFC (vmPFC) in both the learned safety and safety signaling (Eisenberger et al., 2011; Mobbs et al., 2010, Schiller et al., 2008; Suarez-Jimenez et al., 2018).

The present study aimed to investigate the neural mechanisms underlying safety search by manipulating attentional set to safe or dangerous cues using a novel dot-motion paradigm combined with electric shocks. In this paradigm, subjects were asked to judge dot-motion direction while keeping alert to the emergence of safety or danger cues that were associated with either a neutral or aversive outcome (electric shocks). Subjects could be shocked for incorrect responses in the dangerous condition and the shock probability depended on subjects’ performance. Based on the specific role of the vmPFC in safety encoding (Eisenberger et al., 2011; Mobbs et al., 2010; Schiller et al., 2008), we predicted that attentional set to safety search would be mainly under the control of the vmPFC. We additionally predicted that the goal directed attentional network could be involved in searching for dangerous cues (Corbetta and Shulman, 2002; Ptak, 2012; Singh-Curry and Husain, 2009), driven by the higher motivation level occurring in response to cues signaling danger relative to safety (Failing and Theeuwes, 2017; Vuilleumier, 2005).

## Methods and Materials

### Participants

26 healthy students (13 males, mean age = 22.86 years, SD = 4.03) participated in the present study. All subjects were right-handed and had normal or corrected-to-normal vision. None of them reported a history of, or current neurological or psychiatric symptoms. 4 subjects were excluded due to excessive head motion (1 subject), extremely low response accuracy (RA) to shock probability cues (2 subjects) or technical problems during scanning (1 subject). Thus, 22 subjects were included in the final analysis. Written informed consent was obtained from all subjects before study inclusion. The study and all procedures were approved by the local ethical committee and were in accordance with the latest version of the Declaration of Helsinki.

### Stimuli and Procedure

A novel foveal dot-motion paradigm was used in the present study (Figure 1-1), which consisted of 3 threat conditions. The threat condition was indicated by a red (‘dangerous+’: D+), yellow (‘dangerous-’: D-), or green square (safe) for 2 s before each block. Subjects were then asked to complete 2 rating tasks on their confidence and anxiety levels in performing the upcoming blocks using a 1-9 Likert scale within 4 s, followed by a jittered interval of 1-5 s. The foveal moving dots stimuli were generated using Psychopy2 software (v1.83) (Peirce, 2009). Each screen consisted of 100 white random dots (10 × 10 pixels) displayed on a grey background with a moving speed of 0.005 frame. Each dot had a lifetime of 25 frames and was randomly assigned a new position after finishing its lifetime. There were 2 types of tasks in each block and each block comprised 22 trials with a jittered interval of 1-3 s. In the random dots motion (RDM) discrimination task (Figure 1A), dots were presented with a variety of coherence levels ranging from 9%-11%, 14%-16%, and 34%-36% to the left or right direction corresponding to difficult, moderate and easy levels, as determined by an independent pilot study. Dots with these three percentages of coherence moved to left in half of the trials and to the right in the other half. The dots were displayed on the screen for 2 s and subjects were asked to judge the movement direction before they disappeared by pressing either the ‘left’ or ‘right’ response buttons. For incorrect responses, subjects could be punished by one electric shock with an initial probability of 50%, as indicated by a 2-s ‘flash’ symbol feedback. In the shock probability cue detection task (Figure 2A), subjects could minimize the shock probability by giving correct responses to 3 different colored dots corresponding to the 3 threat condition cues. In the D+ condition (red dot), while correct responses did not change the shock probability each incorrect response increased it by 10%. In the D-condition (yellow dot), while each correct response decreased the shock probability by 10%, incorrect responses did not change it. Shock probability changes were indicated by a 2-s feedback. The feedback was a ‘flash’ symbol with specific shock probability changes (e.g., ‘+10%’ or ‘-10%’) appearing above the symbol and the current shock probability below it. In the safe condition (green dot), there was no shock and incorrect responses were given a 2-s ‘cross’ feedback. Only responses with reaction times (RTs) shorter than 650 ms were categorized as correct. The colored dot was presented for 0.5 s and subjects were instructed to respond as fast as possible before it disappeared. These colored dots were carefully counterbalanced in luminance and size. To further show that there was no perceptual bias for the different colored dots, we also conducted a control experiment with an independent sample (N = 18) using a similar dot-motion paradigm, but without administering shock. This showed that there was no significant effect across the different colored dots on either response RTs (ps > 0.196) or RA (ps > 0.210). To maintain maximal continuous attention set for searching of the shock probability cues, subjects were clearly informed that the colored dot could appear at any time point during the 2-s display of white dots on the screen and that only a fast enough correct response would be regarded as a ‘real’ correct response. Following the outcome feedback, subjects were asked to rate their anxiety level while performing the task. In each block, there were 2, 4, or 6 colored dot trials presented in a pseudorandom order among the 22 trials. There were 9 blocks in total with 3 blocks in each threat condition.

**Figure 1.**
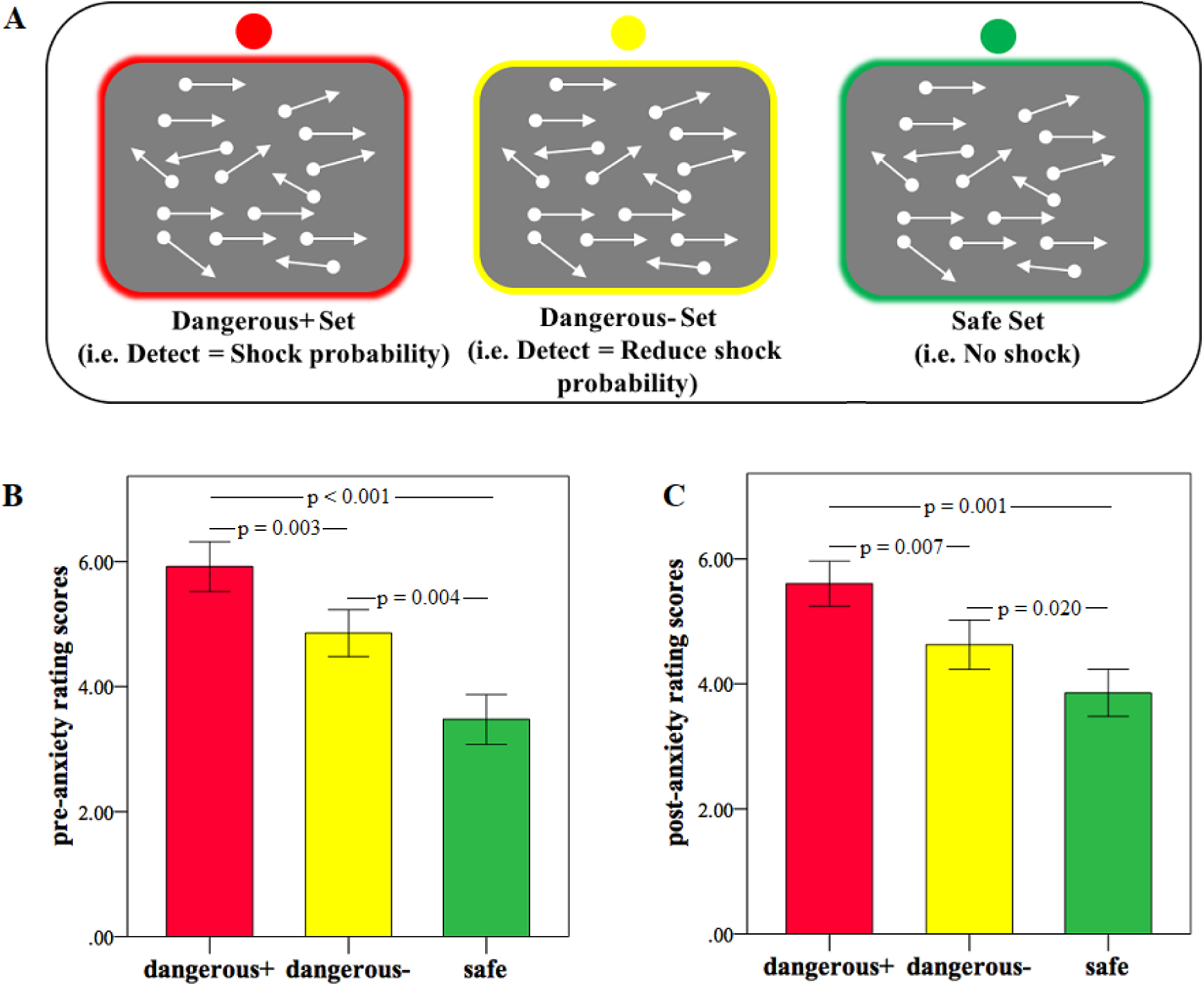
(A) Schematic examples of the random dots motion discrimination task. In this task, subjects were asked to judge the moving direction by pressing the ‘left’ or ‘right’ buttons. (B) Mean pre-anxiety rating scores before each block in each threat condition. (C) Mean post-anxiety rating scores after each block in each threat condition.

**Figure 2.**
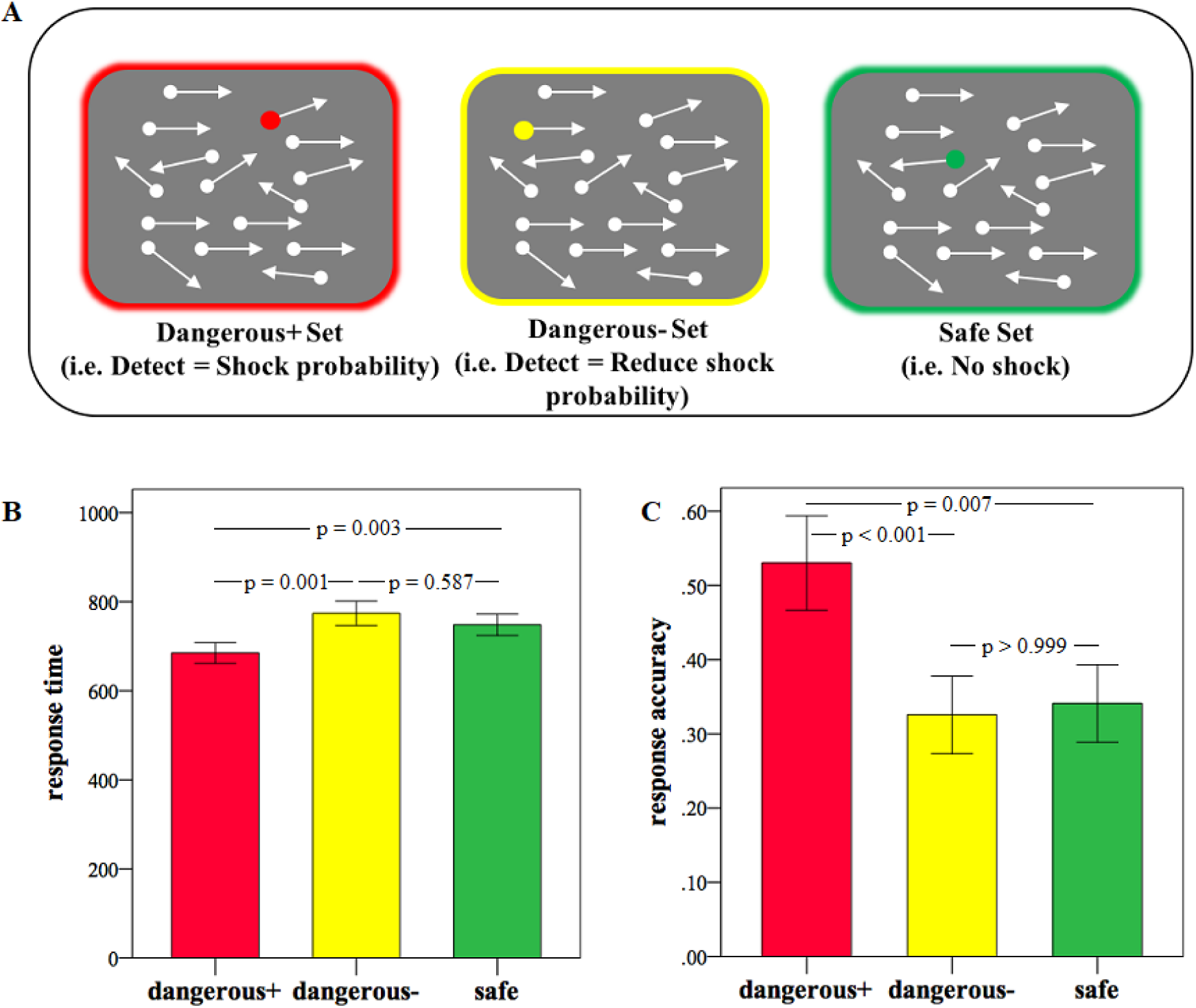
(A) Schematic examples of the cue detection task. Subjects were informed that the colored dot would appear at any time point during the 2-s white dots screen and were instructed to respond as fast as possible before it disappeared. Note that arrows indicate the moving direction of the dots rather than the stimuli *per se*. Mean response time (B) and accuracy (C) to each cue condition in the cue detection task.

### Image Acquisition and Data Analysis

Images were collected using a 3 T, GE Discovery MR750 scanner (General Electric Medical System, Milwaukee, WI, USA). For the fMRI scan, a time series of volumes was acquired using a T2^*^-weighted EPI pulse sequence (repetition time, 2000 ms; echo time, 25 ms; slices, 45; thickness, 3 mm; field of view, 192 × 192 mm; resolution, 64 × 64; flip angle, 77°). High-resolution whole-brain volume T1^*^-weighted images (1 mm isotropic resolution) were acquired obliquely with a three-dimensional spoiled gradient echo pulse sequence before the fMRI scan.

Brain images were processed using the SPM8 software package (Wellcome Department of Cognitive Neurology, London, UK, http://www.fil.ion.ucl.ac.uk/spm/spm8) (Friston *et al*, 1994). The first five images were excluded to achieve magnet-steady images and the remaining functional images were realigned to correct for head motion based on a six-parameter rigid body algorithm. After co-registering the mean functional image and the T1 image, the T1 image was segmented to determine the parameters for normalizing the functional images to Montreal Neurological Institute (MNI) space. Next normalized images were spatially smoothed with an 8 mm full-width at half maximum of Gaussian kernel.

The first-level design matrix included 19 regressors (threat condition cue, confidence rating, pre-anxiety rating, three threat conditions at each difficulty level, three colored threat dots, outcome feedback of direction discrimination task, outcome feedback of colored dots task, delivered shocks, post-anxiety rating) and the 6 head-motion parameters convolved with the canonical hemodynamic response function. On the first level, contrast images for each condition were created for each subject. On the second level, a flexible factorial design was used for the dots moving direction discrimination task to examine the main effect of threat condition and the interaction between threat condition and difficulty. For the cue detection task, a one-way ANOVA within subject design was used to test the main effect of threat condition. For the whole brain analysis, a significance threshold of P < 0.05 false discovery rate (FDR) correction was used with a minimum cluster size of 10 contiguous voxels.

To examine the safety-related effect in a more sensitive way, we further performed a hypothesis-driven region of interest (ROI) analysis in the vmPFC, which has been shown as a core region of safety signaling (Eisenberger et al., 2011; Mobbs et al., 2010; Schiller et al., 2008; Suarez-Jimenez et al., 2018). Furthermore, for the cue detection task, we additionally included ROIs involved in goal-directed attentional processing, namely the intraparietal sulcus (IPS), the inferior parietal lobule (IPL) and frontal eye field, which are core hubs of the frontoparietal attention network (Corbetta and Shulman, 2002; Ptak, 2012; Singh-Curry and Husain, 2009). The vmPFC was derived from the Automated Anatomic Labeling atlas (Tzourio-Mazoyer et al., 2002). The SPL, IPL and IPS were derived from probabilistic maps implemented in Anatomy toolbox 2.1 (Eickhoff et al., 2005). Within these a priori ROIs, a threshold of p < 0.05 family-wise error (FWE) corrected at peak level was set for multiple comparisons.

## Results

### Behavioral results

#### Confidence and anxiety ratings

For ratings, one more subject was excluded due to a data acquisition failure. A repeated-measures ANOVA on the confidence rating scores with threat condition (D+ vs. D-vs. safe) as a within-subject factor revealed a significant main effect (F(2, 40) = 5.91, p = 0.017), with a trend of decreased confidence in the D+ (p = 0.052; mean ± SD = 4.94 ± 1.72) and D-(p = 0.074; 5.49 ± 1.45) conditions relative to the safe condition (6.00 ± 1.48). For the pre-anxiety ratings, there was a significant main effect of threat condition (F(2, 40) = 20.08, p < 0.001). Post-hoc tests showed subjects were most anxious in performing the D+ condition, followed by D‐ and safe conditions (Figure 1B). A significant main effect was also found for the post-anxiety ratings (F(2, 40) = 15.33, p < 0.001), with a similar pattern to the pre-anxiety ratings (Figure 1C).

#### RDM task

We performed a repeated-measures ANOVA on RT and RA with threat condition (D+ vs. D– vs. safe) and difficulty levels (difficult vs. middle vs. easy) as within-subject factors. For RT, this analysis revealed a significant main effect of difficulty (F(2, 42) = 77.40, p < 0.001). Post-hoc analysis reveal that, as expected, subject responded fastest in the easy and slowest in the difficult trials (ps < 0.009; Figure 1–2A). For RA, there was a significant main effect of difficulty (F(2, 42) = 108.08, p < 0.001), with subjects had a higer accuracy in easy than in middle and difficult levels (ps < 0.001; Figure 1–2B). There were no other significant effects (ps > 0.067).

#### Cue detection task

A repeated-measures ANOVA on RT with threat as within-subject factor revealed a significant main effect of threat (F(2, 42) = 11.93, p < 0.001), with faster responses to the D+ cue compared to the D-(p = 0.001) and safe cues (p = 0.003; Figure 2B). For RA, the main effect of threat was also significant (F(2, 42) = 10.78, p < 0.001). Post-hoc test showed that the RA was higher in the D+ cue relative to the D-(p < 0.001) and safe cues (p = 0.007; Figure 2C).

### fMRI results

#### RDM task

In the whole brain analysis, we observed increased activity in left rACC, left dmPFC, right vmPFC, right hippocampus, and bilateral insula in the safe relative to the ‘dangerous +’ conditions (safe > D+; P_FDR_ < 0.05) (Table 1). Comparisons between difficulty levels showed stronger activation in bilateral inferior frontal gyrus, insula, dmPFC/dACC and lingual gyrus and other regions for more difficult than easier tasks (see Table 1-1). However, there were no other siginificant effects in the whole brain analysis (P_FDR_ < 0.05).

**Table 1.** Brain regions activated in safe vs. ‘danger+’ conditions (safe > ‘danger +’).

| Brain Regions | BA | No. Voxels | Peak t-value | x | y | z |
| --- | --- | --- | --- | --- | --- | --- |
| L. Rostral Anterior Cingulate Cortex | 10/32/24 | 1928 | 5.04 | -2 | 36 | -2 |
| Rostral Anterior Cingulate Cortex |  |  | 4.98 | 0 | 38 | 6 |
| Dorsal Medial Prefrontal Cortex |  |  | 4.66 | -10 | 62 | 6 |
| R. Middle Insula | 22/13 | 185 | 4.57 | 46 | 2 | -4 |
| Superior Temporal Gyrus |  |  | 3.98 | 62 | -16 | 2 |
| Middle Temporal Gyrus |  |  | 3.94 | 58 | -8 | -6 |
| L. Middle Insula | 13 | 37 | 4.40 | -36 | 2 | 20 |
| R. Ventral Medial Prefrontal Cortex | 11 | 15 | 3.97 | 4 | 48 | -12 |
| L. Anterior Insula | 13 | 24 | 3.93 | -32 | 10 | -10 |
| R. Hippocampus |  | 14 | 3.69 | 34 | -24 | -10 |
| L. Dorsal Medial Prefrontal Cortex |  | 11 | 3.47 | -8 | -20 | 52 |

All with a $P_{FDR} < 0.05$ corrected threshold and cluster > 10 voxels. MNI coordinates were used. L indicates
left; R indicates right.

The hypothesis-driven ROI analysis further revealed stronger activition in the bilateral vmPFC (left: MNI = −2, 50, −4; t = 4.54; P_FWE_ = 0.001; voxels = 126; Figure 3A; right: MNI = 4, 48, −12; t = 3.97; P_FWE_ = 0.006; voxels = 77) for the safe relative to the D+ conditions (safe > D+). Increased activity was also observed in the left vmPFC (MNI = −8, 52, −12; t = 3.46; P_FWE_ = 0.024; voxels = 4; Figure 3B) in the D-compared to the D+ conditions (D‐ > D+). Given the left vmPFC only has 4 voxels, we further confirmed this effect by extracting the parameter estimates using an independent coordinate (MNI = −6, 51, −15) associated with safety processing in a previous study (Eisenberger et al., 2011). A paired t-test revealed a significantly increased activity of the left vmPFC in the D-(mean = 1.18, SD = 1.79) than the D+ (mean = 0.29, SD = 1.76) conditions (t(21) = 2.91, P = 0.008). Importantly, the left vmPFC overlapped between the ‘safe > D+’ and ‘D‐ > D+’ comparisons (Figure 3C). To exclude the possibilty that the different number of shocks in D+ and D-conditions may confound the findings, we examined whether shock number was associated with the vmPFC activity but found no significant correlations (ps > 0.368).

**Figure 3.**
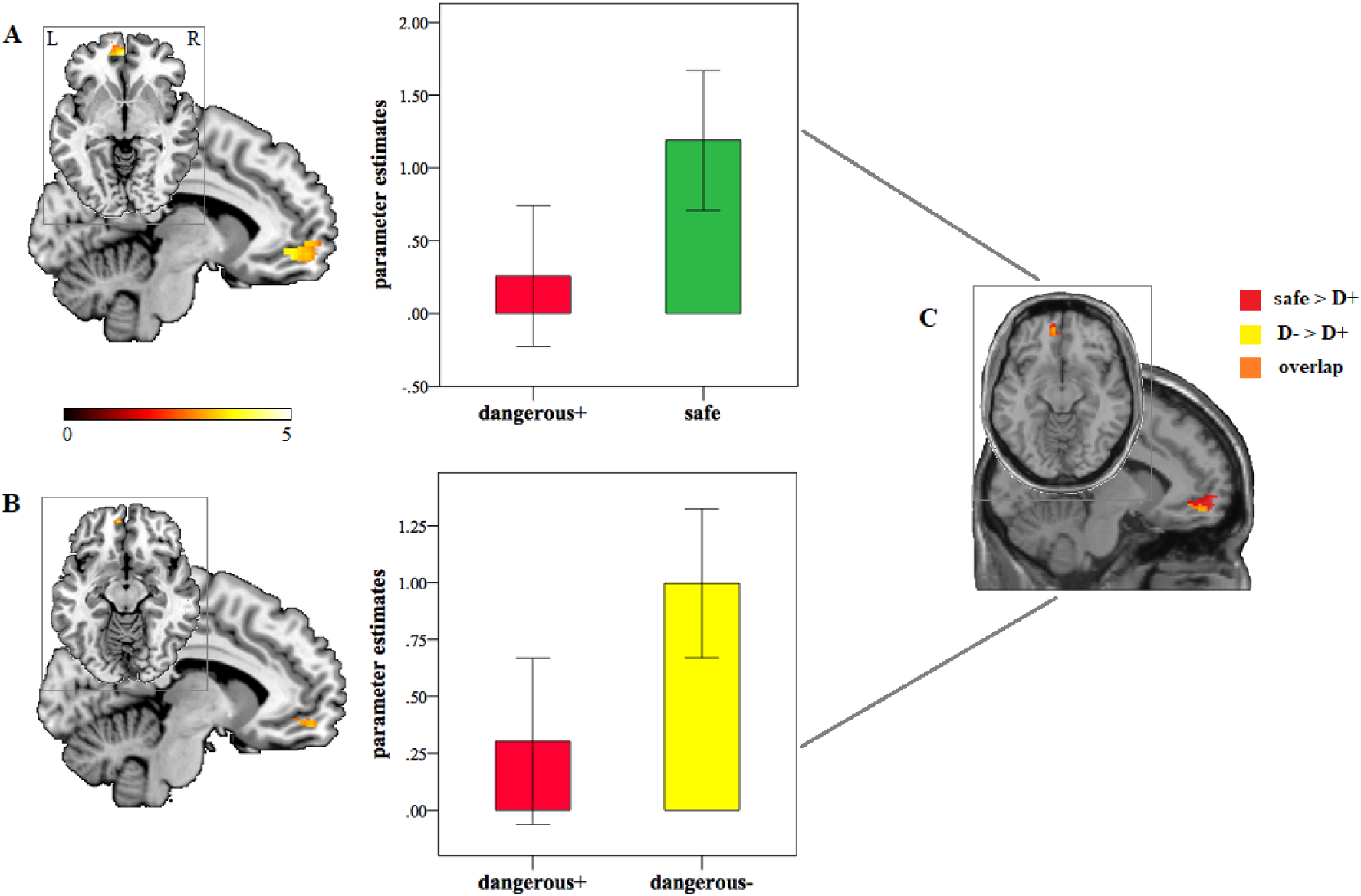
Increased left vmPFC activity in the random dots motion (RDM) task in response to (A) the safe relative to D+ threat conditions (safe > D+) and (B) to the D-relative to D+ threat conditions (D‐ > D+). (C) Overlap between the ‘safe > D+’ and ‘D‐ > D+’ comparisons. Statistic maps were displayed with a P < 0.005 uncorrected threshold. L: left. R: right. D+: ‘dangerous+’. D-: ‘dangerous-’.

#### Cue detection task

The ROI analysis showed stronger activity in the bilateral IPL (left: MNI = −38, −72, 34; t = 4.16; P_FWE_ = 0.044; voxels = 36; right: MNI = 50, −56, 36; t = 4.66; P_FWE_ = 0.013; voxels = 85) and the right IPS (MNI = 44, −52, 34; t = 4.15; P_FWE_ = 0.015; voxels = 13) in response to the D+ compared to the safe cues (D+ > safe cue; Figure 4A). Similar increased activity was also found for the D-relative to the safe cues (D‐ > safe cue) in the right IPL (MNI = 48, −64, 34; t = 4.23; P_FWE_ = 0.038; voxels = 189) and the right IPS (MNI = 46, −50, 44; t = 4.03; P_FWE_ = 0.020; voxels = 33; Figure 4B). For the D‐ vs. D+ comparison (D‐ > D+ cue), we observed a stronger activation in the right IPS (MNI = 44, −38, 48; t = 3.79; P_FWE_ = 0.037; voxels = 10; Figure 4C). To further examine this unpredictable finding, we performed an exploratory psychophysiological interaction (PPI) analysis using the PPI toolbox (McLaren *et al*, 2012) with the right IPS as a seed region (6-mm sphere centered at MNI = 44, −38, 48). Results showed an increased functional connectivity between the right IPS and the left hippocampus (MNI = −30, −30, −12; t = 4.52; P_FWE_ = 0.023; voxels = 14). We also found increased rACC activation (MNI = 2, 34, 4; t = 4.20; P_FWE_ = 0.027; voxels = 39; Figure 4–2) in response to the safe relative to the D+ cues (safe > D+ cue) using a mask from the ‘safe > D+’ contrast (P_FDR_ < 0.05) in the RDM task. There were no significant effects in the whole brain analysis (P_FDR_ < 0.05).

**Figure 4.**
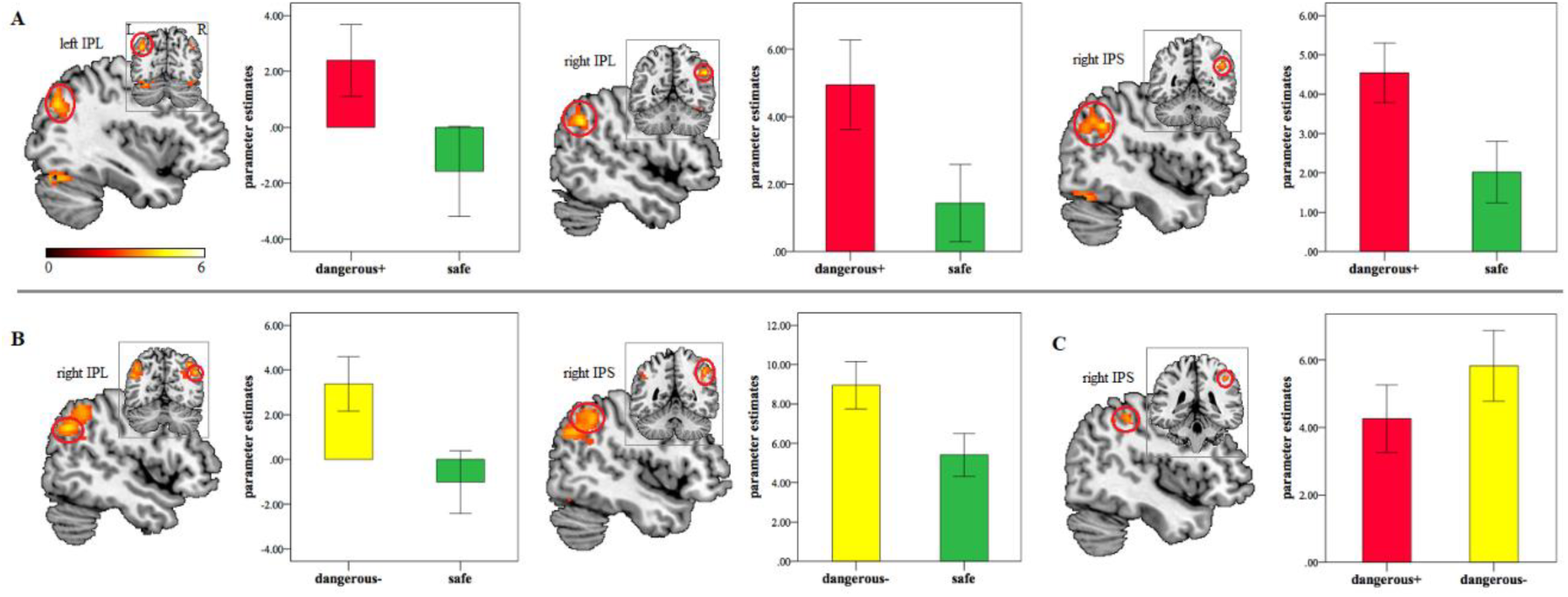
Brain activation in the cues detection task in response to (A) the D+ relative to safe threat cues (D+ > safe), (B) the D-relative to safe cues (D‐ > safe) and (C) the D-relative to D+ cues (D‐ > D+). Statistic maps were displayed with a P < 0.005 uncorrected threshold. IPL: inferior parietal lobule. IPS: intraparietal sulcus. D+: ‘dangerous+’. D-: ‘dangerous-’.

## Discussion

The present study investigated how the human brain is organized to search for safe vs. dangerous cues using a novel RDM paradigm combined with threat of electric shocks. Results showed that subjects tended to be more confident and less anxious in searching the safe relative to the dangerous cues where they could be shocked for incorrect responses. While we found no evidence for significant effects associated with threat conditions on the RDM task performance at the behavioral level, the left vmPFC was recruited both when attention was set to search the safe and D-cues in comparison to the D+ cues. For the cue detection task, while at behavioral level subjects were faster and more accurate in detecting the D+ than the D‐ and safe cues, stronger activity was found in the goal directed attentional networks, including IPL and IPS, for detecting the D+ cues.

Subjects tended to be less confident and more anxious in performing the more threatening relative to the safe searching tasks, which validated the threat manipulation in the present study and coincided with previous findings that the presence of safe cues decreases anxiety to threat (Grillon et al., 1994; Hood et al., 2010). Since shock probability cues were presented in a spatiotemporally random way in the current paradigm, it was beneficial for subjects to respond conservatively to increase accuracy, and thus decrease shocks, and this may have contributed to the absence of behavioral differences across conditions. For the detection of shock probability cues *per se* in the cue detection task, we found similar patterns for RT and RA with subjects responding faster and more accurately to D+ than to D‐ and safe cues. These enhanced behavioral responses could be driven by a higher motivational value of D+ cues, as demonstrated by similar preferential processing of threatening stimuli or more valuable stimuli (such as stimuli associated with higher reward) (Hansen and Hansen, 1988; Hickey et al., 2010).

At the neural level, increased activity was found in the rACC, dmPFC, vmPFC, and hippocampus for the safe relative to the D+ threat conditions in the RDM task. These regions constitute the ‘cognitive fear’ circuitry implicated in elaborate assessment of distal threats and consequently can promote behavioral flexibility and optimize survival decisions (LeDoux and Pine, 2016; McNaughton and Corr, 2004; Price, 2005). Similar neural response patterns have also been evoked by distal threat in previous human fMRI studies (Mobbs et al., 2007, 2009; Qi et al., 2017). Furthermore, the vmPFC was also recruited in the D-compared to the D+ threat conditions and overlapped with the vmPFC identified when attention was set to safe cues. Given the specific role of the vmPFC in learned safety and safety signaling (Eisenberger et al., 2011; Mobbs et al., 2010, Schiller et al., 2008; Suarez-Jimenez et al., 2018), it could be a core region in maintaining attention set not only to safety but also to less inmminent threat. The medial prefrontal areas, especially the vmPFC, receive and integrate inputs from multiple sensory modalities and make them available for higher-level cognitive processes such as emotion regulation, action planning or decision-making (Miller and Cohen, 2001; öngür and Price, 2000; Price, 2005). These regions also connect to the midbrain periaqueductal gray and limbic regions, including the hippocampus, with the former instigating reflexive defensive reactions to imminent threat and the latter involving planning derived from its role in prospective memory (Cohen and O’Reilly, 1996; Miller and Cohen, 2001; Mobbs et al., 2015; Shipley et al., 1991). The vmPFC also coordinates with the hippocampus in mediating fear extinction (Kalisch et al., 2006; Milad et al., 2007). High-level information integration in the medial prefrontal areas would allow more elaborate appraisal of potential threat in the environment and thus generate appropriate levels of anxiety and fear, thereby enabling initiation of optimal actions to threat depending on their perceived imminence. These findings suggest that the neural mechanisms underlying safety search may work in a similar way to how distal threat is encoded and provide the first evidence that the ‘cognitive fear’ circuitry, particularly the medial prefrontal areas, may be specific substrates underpinning attentional set of searching stimuli that are of survival value in the environment, including stimuli signaling safety.

Note that dysfunction of safety processing is also associated with psychiatric disorders (Kong et al., 2014), with panic disorder patients showing impaired learning ability in discriminating between safe and dangerous cues and a less effective fear-reduction by safety cues (Lissek et al., 2009). High trait anxiety individuals also exhibit exaggerated fear generation by safety cues (Haddad et al., 2012) and posttraumatic stress disorder patients fail to inhibit fear response to safety cues (Jovanovic et al., 2010). Thus the medial prefrontal areas could be a target region for potential noninvasive therapeutic interventions, such as real-time fMRI neurofeedback training which has been found to be effective at a clinical level (Watanabe et al., 2017).

Consistent with faster and more accurate behavioral responses, stronger activity in the dorsal frontoparietal attention network, including IPL and IPS, was found for the D+ and D-in comparison to the safe cues in the cue detection task. This suggests an enhanced goal-directed attentional processing for more threatening tasks requiring more cognitive resources. These findings coincide with previous studies showing facilitated processing in the attentional network for higher motivational stimuli such as more threatening or rewarding values (Anderson, 2017; Armony and Dolan, 2002; Failing and Theeuwes, 2017; Vuilleumier, 2005), which has clear adaptive benefits for survival. Note that the D-cues also induced stronger activity in the right IPS than the D+ cues, which could be driven by an increased functional connectivity of the right IPS with the left hippocampus. This is line with the role of hippocampus either in modulating responses to less imminent threat (McNaughton and Corr, 2004; Mobbs et al., 2009; 2015) or in orienting visual attention (Goldfarb et al., 2016; Summerfield et al., 2016). Future studies are necessary to further investigate this question. Furthermore, the rACC was also recruited during processing of the safe compared to the D+ cues. Consistent with its role in attentional set of safety in the RDM task, the enhanced rACC activity may reflect an elaborate evaluation of the utilization of safety, such as a refuge, which is normally associated with protection and opportunity to escape.

Overall, the present study investigated how the human brain encodes safety information by modulating subjects’ attentional set using a novel dot-motion paradigm. Similar to neural mechanisms involved in processing distal threat, the present study demonstrated that attention set of safety mainly recruited medial prefrontal regions of the ‘cognitive fear’ circuitry. Thus, encoding of safety signals may share similar neural substrates with processing of distal threat that allows for flexible threat assessment and consequently increases chances of survival for organisms through exploiting their environment. These findings provide new insights into the role of the medial prefrontal regions in the defensive survival system in encoding stimuli with survival significance.

## Acknowledgements

This study was supported by a grant from the National Natural Science Foundation of China (NSFC, grant number 31700998) to Shuxia Yao and a grant from NARSAD to Dean Mobbs. The authors declare no competing financial interests.

